# A genetic screen identifies O-antigen as essential for *Rickettsia parkeri* survival in macrophages by protecting from inflammasomes and interferon-stimulated genes

**DOI:** 10.64898/2026.06.12.731980

**Authors:** Han Sun, Alejandro A. Guzman, Thomas P. Burke

## Abstract

Lipopolysaccharide (LPS) is highly immunostimulatory, yet it is evolutionarily conserved among many obligate intracellular bacteria for unknown reasons. We report a forward genetic screen to identify factors required for survival of the tick-borne obligate cytosolic pathogen *Rickettsia parkeri* in primary macrophages. The most critical factors were WecA and RmlD, which synthesize O-antigen, the outermost layer of LPS. *wecA* and *rmlD* mutants grew at similar rates to wild type bacteria in epithelial cells, yet in macrophages they were targeted by guanylate binding proteins (GBPs) and they hyperactivated inflammasomes. Survival of O-antigen-deficient mutants was restored >1,000-fold in macrophages lacking Caspases-1 and -11, interferon signaling, and nitric oxide production, suggesting a multifaceted role for O-antigen in protecting against innate immunity. O-antigen was essential for causing disease in mice and protected *R. parkeri* against complement *in vitro*. Despite O-antigen being known as a major target of antibodies, mice immunized with O-antigen-deficient mutants were protected from a lethal rechallenge, suggesting that protection can be elicited independently of O-antigen-targeting antibodies. Together, these findings help resolve a paradox as to why obligate cytosolic bacteria evolutionarily maintain LPS despite it being immunostimulatory, which is that it serves as a multifunctional shield against innate immunity.

**Significance:** Eukaryotic innate immune systems evolved to detect conserved microbial structures as danger signals of infection. Intracellular pathogens, in turn, evolved to hide from innate immunity, yet these mechanisms remain incompletely understood. Here, we performed an unbiased forward genetic screen in macrophages that identified lipopolysaccharide O-antigen as a critical virulence determinant in the tick-borne obligate cytosolic pathogen *Rickettsia parkeri*. We found that O-antigen shields the bacteria from multiple innate immune defenses, including guanylate-binding proteins, inflammasomes, nitric oxide, and complement. These findings reveal why a highly immunostimulatory molecule such as lipopolysaccharide is maintained by an obligate intracellular pathogen and establish O-antigen as a central determinant of *Rickettsia* cytosolic survival with implications for vaccine development.

## Introduction

The *Rickettsia* are a clinically important and diverse genus of arthropod-borne bacteria that obligately reside in the host cell cytosol. Some *Rickettsia* species cause serious human disease including spotted fever and typhus. *Rickettsia parkeri* is one of the spotted fever group (SFG) pathogens, along with the more deadly *R. rickettsii,* which is the causative agent of Rocky Mountain spotted fever. *R. parkeri* causes a moderate form of spotted fever illness in the Americas characterized by fever, rash, and a hallmark eschar skin lesion at the site of tick bite^1,2^. Cases of tick-borne diseases including spotted fever have risen dramatically in recent decades with limited therapeutic options and no approved vaccines^3^, highlighting the need for improved understanding of the molecular mechanisms by which these pathogens cause disease.

Immune cells play a critical role in *Rickettsia* infections, and a variety of studies suggest that they are the dominant cell types targeted by *R. parkeri.* Skin biopsies of human infections found that *R. parkeri* is primarily associated with immune cells, including macrophages, neutrophils, and monocytes^2,4^. Similar observations of immune cell targeting were made during *R. parkeri* infection of non-human primates when delivered by ticks^5^, and upon experimental infection of guinea pigs^6^. We recently reported that in wild type (WT) mice, which resist infection, and double knockout mice deficient in interferon-I and -II receptors (*Ifnar^-/-^Ifngr^-/-^*) that are susceptible to eschar-associated rickettsiosis and succumb to infection, *R. parkeri* is mainly associated with immune cells in the skin, as well as in internal organs including the spleen, liver, and lung upon dissemination^7^. After adherence and uptake in immune cells or other cell types, *Rickettsia* quickly escape to the host cytosol, where they replicate and spread from cell-to-cell using actin-based motility^8^. As the eukaryotic cytosol is rich in host defense factors, *Rickettsia* must presumably evade innate immune detection and killing by host defenses. *Rickettsia* have therefore likely evolved sophisticated mechanisms to evade innate immunity and survive in immune cells; however, such factors remain largely unknown.

Due to their obligate intracellular nature, laborious approaches are needed for genetic manipulation, and thus *Rickettsia* virulence factors are largely enigmatic as compared to facultative pathogens. Few forward genetic screens have been reported to identify *Rickettsia* virulence factors through unbiased approaches^9^ and only a handful of secreted effectors and surface factors have been described in detail for their roles in promoting *R. parkeri* pathogenesis^10–12^. One such factor is the outer membrane protein B (OmpB), which is required for *R. parkeri* to evade antibacterial autophagy and survive in macrophages^13^. Mutants lacking *ompB* were highly attenuated in mice, suggesting that survival in macrophages dictates pathogenicity *in vivo*. This concept has also been suggested upon comparisons of how non-pathogenic versus pathogenic *Rickettsia* survive in macrophage-like cells^14^. Thus, survival in immune cells may be a strong correlate for rickettsial virulence *in vivo*.

The *Rickettsia* outer membrane contains lipopolysaccharide (LPS), which is composed of lipid A, core oligosaccharides, and outward-facing polysaccharides known as the O-antigen. *Rickettsia* LPS is similar in structure to other Gram-negative bacteria^15,16^. Rickettsial LPS is sensed by TLR4, which elicits cytokine secretion and protection in mice^17,18^. Moreover, cells lacking the non-canonical inflammasome Caspase-11, which detects LPS, undergo reduced cell death upon infection with *R. parkeri* and other *Rickettsia,* further suggesting that *Rickettsia* LPS is immunostimulatory^19,20^. Considering that some facultative pathogens like *Francisella* species evolved mechanisms to modify their LPS and evade immune stimulation^21^, it is paradoxical that LPS is so highly conserved among SFG *Rickettsia*, as these microbes inhabit the eukaryotic environment in both mammals and ticks for their entire lifecycle.

To identify the *Rickettsia* virulence factors required for survival in immune cells, we performed a forward genetic screen for *R. parkeri* survival in primary macrophages. We found that O-antigen is critical for survival in these cells, but not for growth in epithelial cells or in immortalized macrophages. Mechanistically, O-antigen enabled *R. parkeri* to avoid inflammasomes and anti-rickettsial ISGs. *In vivo,* we found that O-antigen was crucial for causing disease in mice but surprisingly, that protective immunity is conferred by mutants lacking O-antigen. Through the lens of evolution, these findings provide an explanation for why obligate cytosolic bacteria maintain LPS despite its potent immunostimulatory properties, as it is part of a multilayered surface defense system that enables *R. parkeri* to evade innate immune host defenses.

## Results

### A forward genetic screen identifies O-antigen biosynthesis factors required for *R. parkeri* survival in primary macrophages

*R. parkeri* targets immune cells including macrophages *in vivo,* yet the virulence factors required for survival in these cells remain largely unknown. To identify *R. parkeri* factors required for survival in macrophages, we screened previously generated libraries of *R. parkeri* transposon mutants^12,22^ for growth in murine bone marrow-derived macrophages (BMDMs), which are permissive for growth of WT bacteria^19^. BMDMs were infected with each mutant alongside wild type (WT) *R. parkeri* as a control and plaque forming units (PFUs) were measured at 2 and 72 hpi. After screening 320 mutants, we observed that the majority of bacteria grew at rates similar to WT *R. parkeri* (**Fig. 1A**). PFU at 72 h were divided by the PFU at 2 h to calculate the growth rate, revealing four mutants with a growth rate below 1, meaning that bacterial numbers declined from 2 to 72 h in BMDMs (**Fig. 1B**). Two of the four were insertions in *ompB,* which were previously reported to suffer in BMDMs due to restriction by antibacterial autophagy^13^. The screening data also aligned with the previous report on mutants lacking protein methyltransferases 1 and 2 (PkmT1 and PkmT2), which had a reduced ability to grow in BMDMs that was to a lesser extent than WT bacteria but was not below a growth rate of 1^12^. The two most severely restricted mutants contained insertions in *MC1_07070* (locus tag MC1_RS06510) and *MC1_02580* (locus tag MC1_RS02345), which were annotated as a UDP-N-acetylglucosaminyltransferase and dTDP-4-dehydrorhamnose reductase, respectively.

**Figure 1:**
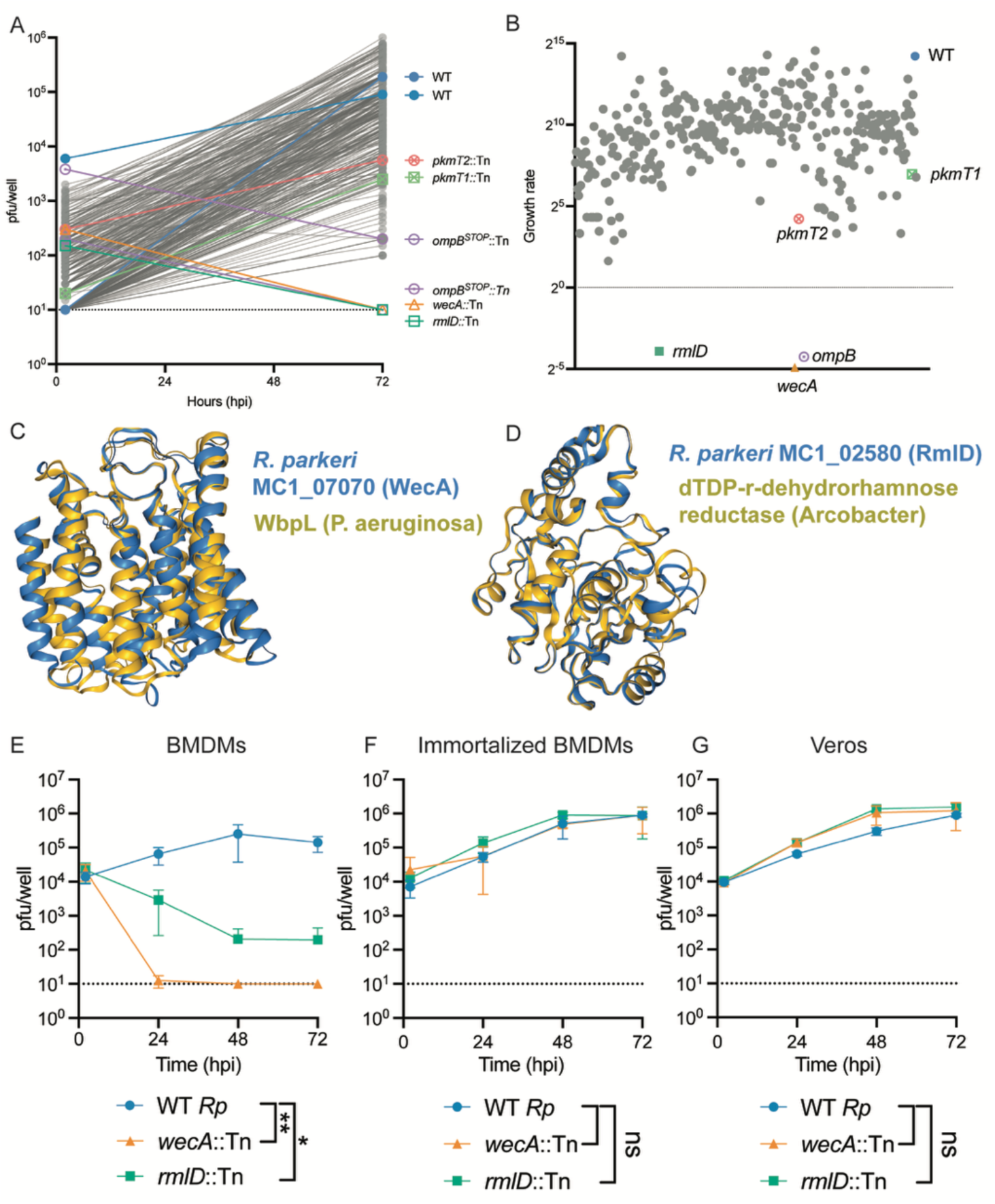
A forward genetic screen identifies O-antigen as required for *R. parkeri* survival in macrophages. **A, B)** Bone marrow-derived macrophages (BMDMs) were infected with 320 different *R. parkeri* mutants and measured for plaque forming units (PFUs) at 2 and 72 hours postinfection. Growth is shown in (A) and rate (PFU at 72 hpi over PFU at 2 hpi) is shown in panel B. **C)** A structural prediction of *R. parkeri* protein WecA (MC1_07070) as determined by AlphaFold was used to identify structural homologs with FoldSeek. The highest similarity to the AFDB-PROTEOME database was glycosyltransferase WbpL from Pseudomonas aeruginosa PAO1 (TM-score 0.84, RMSD: 3.56, E-Value 1.63e-16), as shown. **D)** A structural prediction of *R. parkeri* protein RmlD (MC1_02580) as determined by AlphaFold was used to identify structural homologs with FoldSeek. The highest similarity to the AFDB50 database was dTDP-4-dehydrorhamnose reductase from *Arcobacter* Sp. CECT 8986 (TM-score: 0.97, RMSD: 0.96, E-value 2.54e-43.), as shown. **E)** Abundance of WT *R. parkeri* (n = 3), *wecA*::Tn (n = 4), and *rmlD*::Tn (n = 5) in BMDMs infected at a MOI of 1. **F)** Abundance of WT *R. parkeri* (n = 6), *wecA*::Tn (n = 6), and *rmlD*::Tn (n = 6) in immortalized BMDMs (L3OGs) infected at a MOI of 1. **G)** Abundance of WT *R. parkeri* (n = 4), *wecA*::Tn (n = 6), and *rmlD*::Tn (n = 6) in Vero cells infected at a MOI of 1. Infected cells were lysed at indicated timepoints for plaque assay. Dotted line indicates the limit of detection. Data are expressed as means ± SD. Statistics were performed with a two-way ANOVA, *p<0.05, **p<0.01, ns = not significant.

We focused our attention on MC1_07070 and MC1_02580 due to their severe restriction observed in the screen. These mutants were also previously identified in a genetic screen for *R. parkeri* mutants that are ubiquitylated, where Engström and colleagues found that these factors were required for O-antigen biosynthesis in *R. parkeri* as determined by using an O-antigen targeting antibody^12^. MC1_07070 was annotated as a UDP-N-acetylglucosaminyltransferase and BLAST results showed the closest homologs to be MraY glycosyltransferases. We sought to better characterize if MC1_07070 was structurally similar to O-antigen biosynthesis factors and determined a predicted structure with AlphaFold (**Fig. 1C**). Using FoldSeek, we then identified structures of proteins with known and validated functions, which revealed that MC1_07070 had the highest similarity to the glycosyltransferase WbpL from Pseudomonas aeruginosa PAO1 (**Fig. 1D**). WbpL was determined biochemically to be required for *Pseudomonas* O-antigen biosynthesis^23^, aligning with the report from Engström *et al* that this factor is an O-antigen biosynthesis factor. Regarding MC1_02580, which is annotated as a dTDP-4-dehydrorhamnose reductase, such proteins are widely appreciated as being required for O-antigen biosynthesis, and we found high similarity between the *R. parkeri* homolog and such reductases in other species (**Fig. 1D**), including *Shigella* and *Salmonella*, where this protein’s activity has been determined biochemically^24^. As these factors were previously referred to as WecA and RmlD in *R. parkeri* according to their functional relatives in *Shigella* and other species^25^, we will refer to them here as WecA (MC1_07070) and RmlD (MC1_02580). Based on the structural similarities to homologs of these proteins in other species and the lack of any detectable O-antigen of these mutants in *R. parkeri,* this strongly suggests that they are responsible for the first and second steps of O-antigen biosynthesis in *R. parkeri*.

As the initial screen only used early (3 h) and late (72 h) time points, we sought to better define the kinetics of *wecA*::Tn and *rmlD*::Tn restriction in BMDMs. Upon measuring PFUs every 24 h, we found that the *wecA* and *rmlD* mutants were dramatically restricted in BMDMs as soon as 24 hpi. When compared to other *R. parkeri* mutants that were previously reported as attenuated *in vivo,* the O-antigen-deficient mutants were more restricted than *pkmT1*, *pkmT2*, or *ompB* mutants in BMDMs (**Fig. 1E**)^12,13^. Interestingly, the restriction was specific to primary macrophages, as both *wecA* and *rmlD* mutants grew robustly in an immortalized BMDMs (**Fig. 1F**) and in Vero cells (**Fig. 1G**). Although many previous reports on *Rickettsia* virulence factors use THP-1 or other macrophage cell lines, this finding emphasizes the importance of evaluating *Rickettsia* virulence factors in primary cells, as these mutants would not have been identified using immortalized macrophages. Together, these data revealed that O-antigen is critical for *R. parkeri* survival in primary macrophages.

### Autophagy is not a major host defense restricting O-antigen-deficient mutants in macrophages

It was unclear how O-antigen promoted *R. parkeri* intracellular survival in macrophages. WecA and RmlD were previously reported to protect *R. parkeri* from ubiquitin, albeit to a lesser extent than OmpB, PkmT1, and PkmT2^12,13^. *R. parkeri ompB* mutants were restricted for survival in WT BMDMs, and their growth was restored in BMDMs from mice lacking the autophagy component Atg5^13^. We therefore hypothesized that survival of *wecA* and *rmlD* mutants would be restored in *Atg5^-/-^* BMDMs. To test this hypothesis, we measured survival of these mutant bacteria in BMDMs generated from conditional mutant mice where *Atg5* was excised (LysM-Cre mice crossed to *Atg5^loxP^* mice) and control cells and compared survival of the mutant bacteria to WT *R. parkeri*. Upon measuring bacterial burdens over time, *wecA*::Tn and *rmlD*::Tn survival was not restored in *Atg5-*deficient cells (**Supplemental Fig. 1A**), suggesting that *R. parkeri* O-antigen protects against distinct innate immune factors as OmpB.

### O-antigen enables *R. parkeri* to evade restriction by guanylate-binding proteins (GBPs)

*R. parkeri* are unusual among other intracellular bacterial pathogens in that they are strongly restricted by type I interferons (IFN-I), and we previously found that sub-populations of *R. parkeri* could be targeted and killed by interferon-stimulated genes (ISGs) including guanylate binding proteins (GBPs)^19^. GBPs can localize to intracellular bacteria and cause the release of LPS, activating the Caspase-11 inflammasome^26^. We therefore hypothesized that O-antigen was required for *R. parkeri* to evade killing by GBPs. To test this hypothesis, we infected BMDMs and measured release of lactate dehydrogenase (LDH). LDH is a host cytosolic enzyme that is released to the supernatants upon pyroptosis and thus serves as a readout for host cell inflammasome activation and pyroptosis. At a multiplicity of infection (MOI) of 1, WT cells elicited 6% LDH release at 6 hpi and 40% LDH release by 24 hpi, similar to previous findings^19^. In contrast, *wecA* and *rmlD* mutants caused significantly more cell death at both times, 35% and 60%, respectively (**Fig. 2A**, **2B**). To determine if this killing was inflammasome-dependent, we infected BMDMs lacking Caspase-1 and -11, and indeed the cell death from WT, *wecA*::Tn, and *rmlD*::Tn *R. parkeri* was dramatically reduced (**Fig. 2A**, **2B**). To determine if the increased cell death was due to killing by GBPs, we infected BMDMs lacking GBPs on chromosome 3 (*Gbp^chr3^*^-/-^). Cell death from *wecA* and *rmlD* mutants was reduced from 40% to 4% at 6 hpi (**Fig. 2A**), suggesting a critical role for O-antigen in protecting against GBPs at early time points postinfection. At 24 hpi, LDH release in *Gbp^chr3^*^-/-^ cells was also significantly reduced upon infection with *rmlD*::Tn, albeit to a lesser extent, while *wecA*::Tn had similar LDH release as in WT cells (**Fig. 2B**). These findings suggest an important role for *R. parkeri* O-antigen in protecting against GBP-mediated pyroptosis. Notably, these findings contrast the paradigm in the field^27^ established with *Shigella,* whereby O-antigen-deficient mutants have decreased targeting of GBPs^28^.

**Figure 2:**
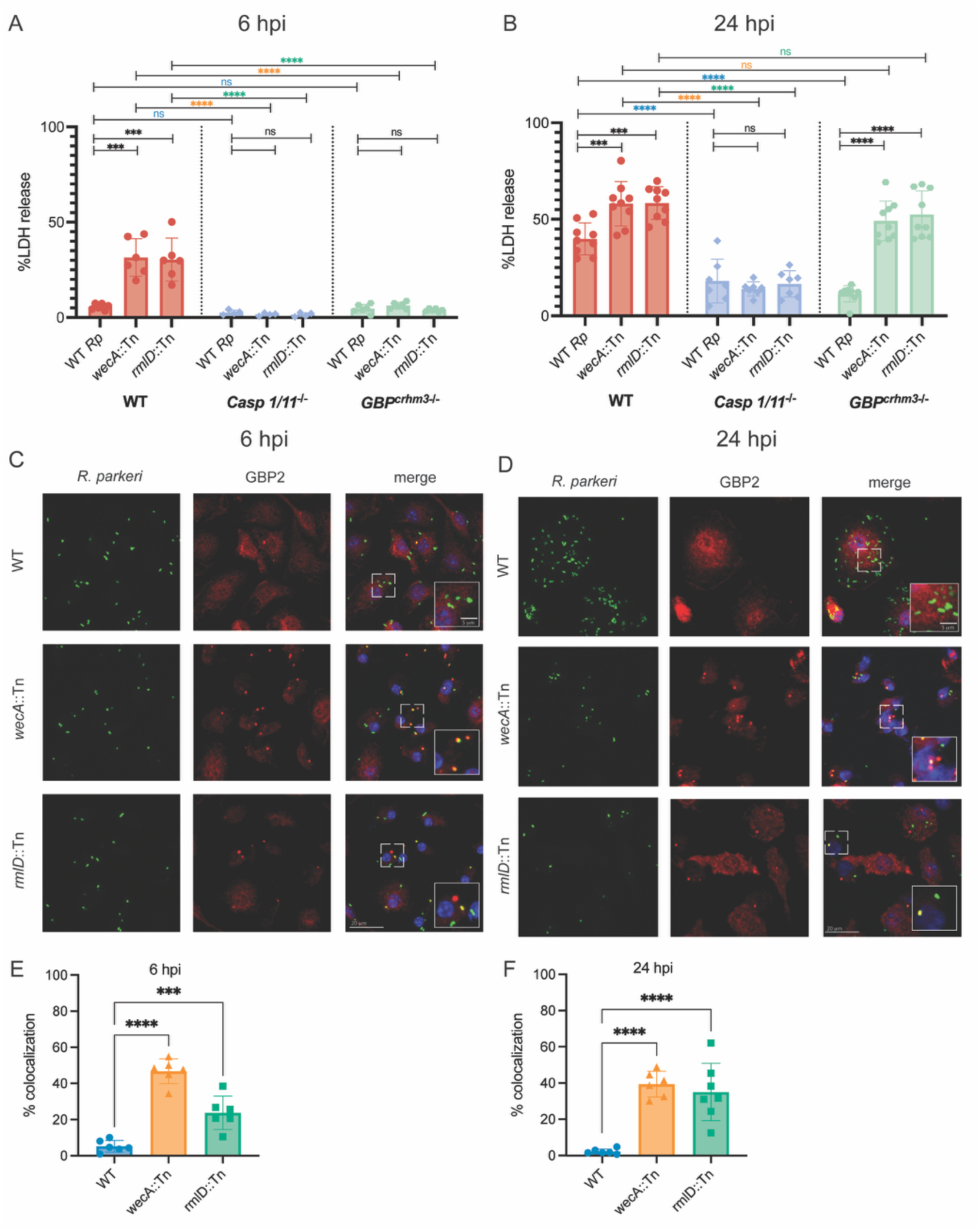
O-antigen enables *R. parkeri* to evade restriction by guanylate-binding proteins (GBPs). **A, B)** Quantification of host cell death during WT, *wecA*::Tn and *rmlD*::Tn *R. parkeri* infection of WT, *Casp1/11^-/-^*, and *Gbp^chr3-/-^* BMDMs at 6 hours **(A)** and 24 hours **(B)** postinfection. For 6 hpi, WT (n = 6), *Casp1/11^-/-^* (n = 4), and *Gbp^chr3-/-^* (n = 6). For 24 hpi, WT (n = 9), *Casp1/11^-/-^* (n = 7), and *Gbp^chr3-/-^* (n = 9). Statistics were performed with a one-way ANOVA. **C, D)** Representative images of BMDMs infected with WT, *wecA*::Tn and *rmlD*::Tn *R. parkeri* at a MOI of 1 and imaged at 6 hpi **(C)** and 24 hpi **(D)**. BMDMs were pretreated with IFN-β overnight prior to infection. Coverslips were stained with an α-GBP2 antibody (red) and an α-*Rickettsia* antibody (green). Scale bar = 20 µm. **E, F)** Quantification of percent colocalization of *R. parkeri* and GBP2, from images in **(C,D)**. Quantification is an average of at least 3 separate experiments totaling >300 bacteria. Data are expressed as means + SD and were compared with a one-way ANOVA, ***p<0.001, ****p<0.0001, ns = not significant.

To determine whether the GBPs directly targeted O-antigen-deficient mutants, we employed immunofluorescence microscopy and measured colocalization of GBP2 with the bacteria in the presence of IFN-β. Similar to previous findings^19^, 7% of WT *R. parkeri* colocalized with GBP2 at 6 hpi and 3% colocalized at 24 hpi (**Fig. 2C**–**2F**). In contrast, *wecA* mutants had 35-40% positive colocalization while *rmlD* mutants had 35 and 45% colocalization at 6 and 24 hpi, respectively (**Fig. 2C**–**2F**), demonstrating that O-antigen protected the *R. parkeri* surface from targeting by GBP2. In summary, these findings demonstrate that O-antigen protects *R. parkeri* from GBP-mediated targeting and GBP-mediated inflammasome activation.

### O-antigen protects *R. parkeri* from nitric oxide-mediated killing

We next sought to determine whether O-antigen protected *R. parkeri* from other ISGs including nitric oxide (NO), which can restrict bacteria through a variety of mechanisms, including damaging the bacterial membrane, DNA, and proteins, although it remains unknown if O-antigen can protect intracellular bacteria from NO-mediated killing. To determine if O-antigen protects *R. parkeri* from NO-mediated killing and inflammasome activation, BMDMs were treated with L-NIL, a specific iNOS inhibitor, infected with WT, *wecA*::*Tn*, or *rmlD*::Tn *R. parkeri,* and LDH release was measured. Inhibiting iNOS did not significantly reduce LDH release (**Fig. 3A**), suggesting that NO did not play a major role causing cell death to elicit inflammasome activation. To determine the relative roles for GBPs versus iNOS, *Gbp^chr3-/-^* cells were treated with L-NIL, infected, and LDH release was measured. GBP deficiency dramatically reduced LDH release at 6 hpi, while inhibiting iNOS did not further reduce LDH release at 24 h (**Fig. 3B**). In addition to these infection conditions, we also sought to more directly examine the effects of NO on *R. parkeri*, and so we performed an axenic experiment where the bacteria were incubated with NO in liquid media, which was reported to restrict *R. rickettsii* and *R. parkeri* in a dose-dependent manner^29,30^. The *wecA* and *rmlD* mutants were treated with NONOate, a spontaneous NO chemical donor, alongside a WT *R. parkeri* control in liquid media. Over time the bacteria were then added to confluent Vero cells and PFU were measured. As expected, NONOate restricted WT *R. parkeri* in a dose-dependent manner (**Fig. 3C**). The O-antigen-deficient mutants were more severely restricted than WT, as *wecA* mutants were >32-fold more restricted than WT, while the *rmlD* mutants were >13-fold more restricted than WT (**Fig. 3C**). Since NO-mediated restriction of WT *R. rickettsii* was previously reported to be rescued by the addition of exogenous ATP^29^, we then asked whether O-antigen mediated protection from NO was redundant with NO-mediated depletion of ATP. WT and O-antigen-deficient mutants were treated with NONOate in the presence or absence of ATP. The addition of ATP still rescued killing of the O-antigen-deficient mutants by NO (**Fig. 3D**), suggesting that O-antigen-mediated protection is a distinct layer of protection from the role of NO on altering *Rickettsia* metabolic activity. Together, these findings suggest that O-antigen plays a role in protecting *R. parkeri* from NO.

**Figure 3:**
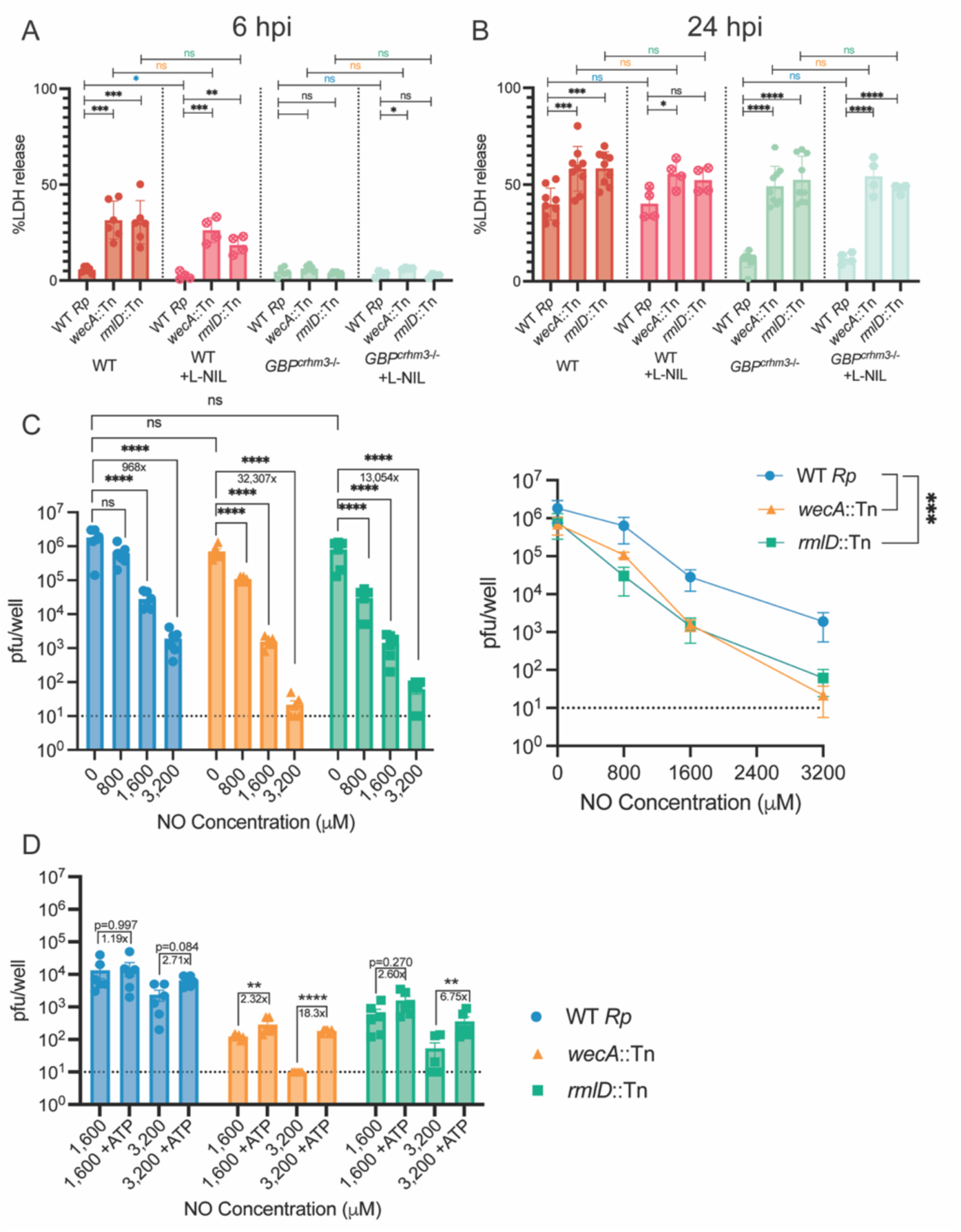
O-antigen helps protect *R. parkeri* from nitric oxide. **A, B)** Quantification of host cell death as measured by LDH release into supernatants during WT, *wecA*::Tn and *rmlD*::Tn *R. parkeri* infection of WT and *Gbp^chr3-/-^* BMDMs at 6 hours **(A)** and 24 hours **(B)** postinfection (n = 6). 1 mM L-NIL was added at indicated conditions (n = 4) 1 hpi. Statistics were performed with a one-way ANOVA. **C)** Abundance of WT, *wecA*::Tn, and *rmlD*::Tn incubated with increasing concentrations of NONOate after 30 minutes. Dotted line represents limit of detection. Data are a combination of six individual experiments and are expressed as means ± SD. Statistics were performed with a lognormal one-way ANOVA (left) and a two-way ANOVA (right). **D)** Abundance of WT, *wecA*::Tn, and *rmlD*::Tn incubated with increasing concentrations of NONOate with the addition of 1 mM ATP after 1 h. Dotted line represents the limit of detection. Data are a combination of six individual experiments and are expressed as means ± SD. Statistics were performed with a lognormal one-way ANOVA. *p<0.05, **p<0.01, ***p<0.001, ****p<0.0001, ns = not significant.

### O-antigen protects *R. parkeri* from inflammasomes and ISGs to survive in macrophages

We next sought to better define the role for O-antigen in protecting against inflammasomes. To determine which inflammasomes were hyperactivated by O-antigen-deficient *R. parkeri* mutants, we infected BMDMs lacking the individual components required for inflammasome-mediated cell death. We hypothesized that pyroptosis was perhaps initiated by the AIM2 inflammasome, which senses cytosolic DNA and protects against O-antigen mutant *Francisella* species^31^, or by NLRP3, which has numerous activators. However, in both *Aim2^-/-^* and *Nlrp3^-/-^*cells, the *wecA* and *rmlD* mutants still elicited significantly increased cell death as compared to the WT BMDM control (**Fig. 4A**). As LDH release was lost in double mutant *Casp1/11*^-/-^ cells, we next sought to determine which caspase was more critical. Curiously, we also observed similar cell death between WT cells and cells lacking either Caspase-1 (*Casp1^-/-^*) or Caspase-11 (*Casp11*^-/-^) (**Fig. 4B**), suggesting that these pathways compensate for one another, and we therefore next evaluated other markers of inflammasome activation.

**Figure 4:**
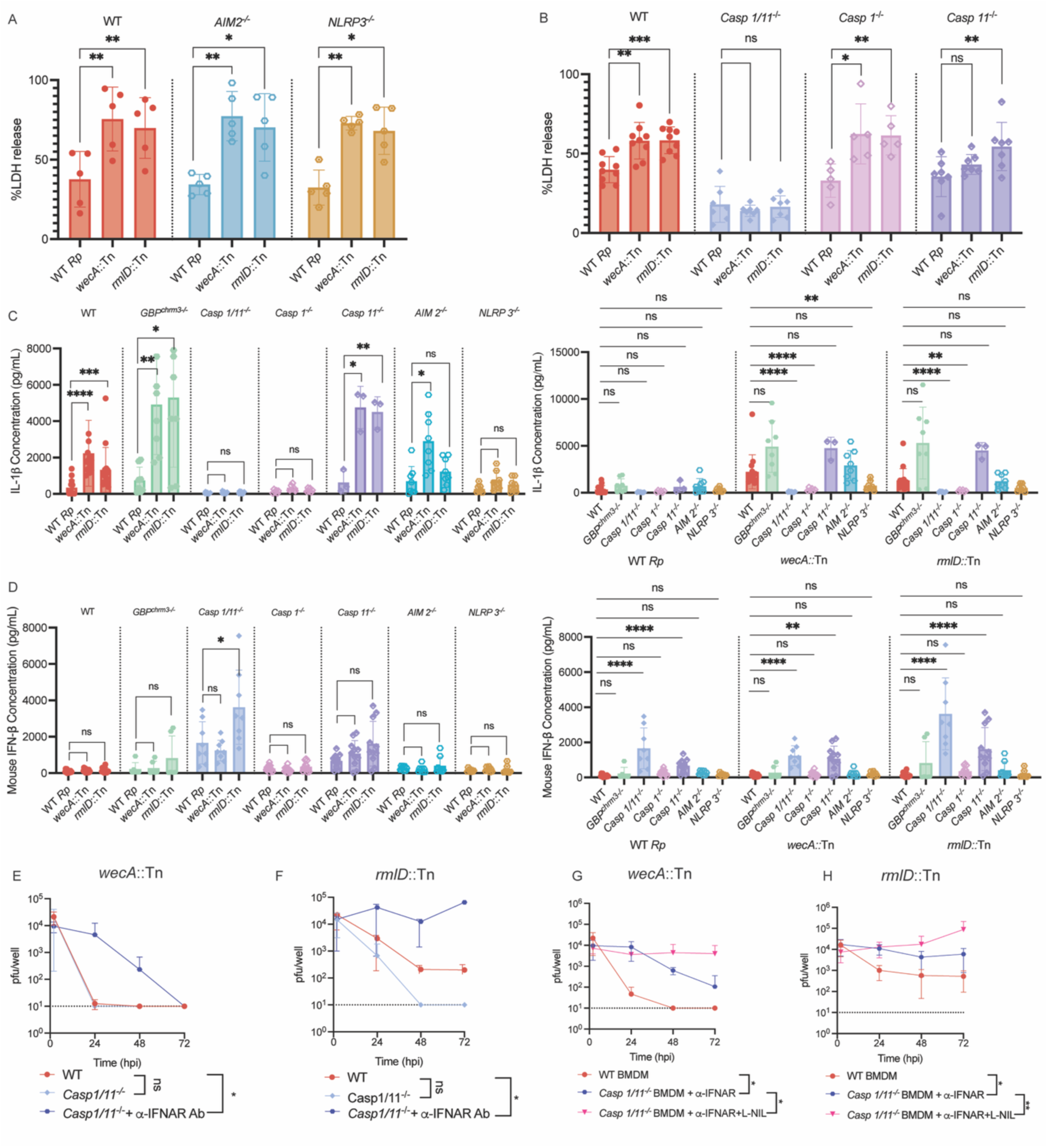
O-antigen protects *R. parkeri* from inflammasome activation in macrophages. **A)** Quantification of host cell death as measured by LDH release into supernatants during WT, *wecA*::Tn and *rmlD*::Tn *R. parkeri* infection of WT, *Aim2^-/-^,* and *Nlrp3^-/-^* BMDMs (n= 5). **B)** Quantification of host cell death during WT, *wecA*::Tn and *rmlD*::Tn *R. parkeri* infection of WT (n = 9), *Casp1/11^-/-^* (n = 7)*, Casp1^-/-^* (n = 5), and *Casp11^-/-^* BMDMs (n = 7). LDH release was measured at 24 hpi after *R. parkeri* infection at a MOI of 1. Data are expressed as means ± SD. Statistics were performed with a one-way ANOVA. **C)** Quantification of IL-1β concentration during WT, *wecA*::Tn and *rmlD*::Tn *R. parkeri* infection of WT (n = 15), *Gbp^chr3-/-^* (n = 8), *Casp1/11^-/-^* (n = 11)*, Casp1^-/-^* (n = 6)*, Casp11^-/-^* (n = 3)*, Aim2^-/-^* (n = 8), and *Nlrp3^-/-^* (n = 7) BMDMs. IL-1β concentration was measured at 24 hpi after *R. parkeri* infection at a MOI of 1. Data are expressed as means ± SD. Statistics were performed with a lognormal one-way ANOVA. **D)** Quantification of IFN-β concentration during WT, *wecA*::Tn and *rmlD*::Tn *R. parkeri* infection of WT (n = 13), *Gbp^chr3-/-^* (n = 6), *Casp1/11^-/-^* (n = 8)*, Casp1^-/-^* (n = 12)*, Casp11^-/-^* (n = 11)*, Aim2^-/-^* (n = 8), and *Nlrp3^-/-^* (n = 8) BMDMs. IFN-β concentration was measured at 24 hpi after *R. parkeri* infection at a MOI of 1. Data are expressed as means ± SD. Statistics were performed with a lognormal one-way ANOVA. **E, F)** Abundance of *wecA*::Tn (n = 4) and *rmlD*::Tn (n = 5) in WT, *Casp1/11^-/-^*, and *Casp1/11^-/-^* BMDMs supplemented with 1 mM α-IFNAR antibody 1 hpi. BMDMs were infected at a MOI of 1. **G, H)** Abundance of *wecA*::Tn and *rmlD*::Tn in WT (n = 10), *Casp1/11^-/-^* BMDMs supplemented with 1 mM α-IFNAR antibody 1 hpi (n = 10), and *Casp1/11^-/-^* BMDMs supplemented with both 1 mM α-IFNAR antibody and 1 mM L-NIL 1 hpi (n = 6). BMDMs were infected at a MOI of 1. Dotted line represents the limit of detection. Data are expressed as means ± SD. Statistics were performed with a two-way ANOVA. *p<0.05, **p<0.01, ***p<0.001, ****p<0.0001, ns = not significant.

Apart from host cell death, inflammasome activation causes IL-1β processing and release, which requires Caspase-1 and NLRP3. To determine if O-antigen masked *R. parkeri* from detection and IL-1 β release, we measured IL-1β secretion into supernatants upon infection of mutant BMDMs lacking inflammasome components. In WT cells, *wecA* and *rmlD* mutants caused dramatically increased secretion of IL-1β, confirming the LDH results that O-antigen protects *R. parkeri* from inflammasome activation (**Fig. 4C**). As expected, IL-1β secretion was ablated in cells lacking *Casp1/11*, *Casp1*, or *NLRP3* (**Fig. 4C**). IL-1β release was similar between WT and *Aim2*^-/-^ cells, suggesting no role for AIM2 in this process. IL-1β was not reduced in *Casp11^-/-^* cells, and was in fact increased, supporting the notion that Caspase-1 is compensating for the loss of Caspase-11 in these cells.

Upon GBP-mediated killing of *R. parkeri,* inflammasome deficient cells hyperproduce IFN-β via cGAS^19^, and so we measured IFN-β production in WT and inflammasome-deficient cells as a proxy for bacteriolysis. WT *R. parkeri* elicited increased IFN-β production in *Casp1/11^-/-^* or *Casp11^-/-^*cells. In WT BMDMs, *wecA* and *rmlD* mutants led to similar amounts of IFN-β secretion as WT *R. parkeri* (**Fig. 4D**). *wecA* and *rmlD* mutants also had increased secretion of IFN-β in *Casp1/11^-/-^* and *Casp11^-/-^* cells. Significantly more IFN-β was produced upon infection of *Casp1/11*^-/-^ cells with the *rmlD* mutant, however, the responses otherwise were largely similar to one another. These findings are consistent with Caspase-11-mediated inflammasome activation limiting IFN-β production.

We next sought to determine whether the inflammasome and IFN-β responses were responsible for restricting O-antigen-deficient mutants and measured bacterial abundance over time in *Casp1/11^-/-^* cells. To account for the increased IFN-β produced in inflammasome-deficient cells, these experiments were performed in parallel using cells treated with an α-IFNAR antibody to block IFN-β signaling. *wecA*::Tn *R. parkeri* did not increase in bacterial abundance in untreated *Casp1/11*^-/-^ BMDMs, but interestingly, was increased >1,000-fold in cells treated with the α-IFNAR antibody at 24 and 48 hpi (**Fig. 4E**). The *rmlD* mutant had a similar restriction in *Casp1/11^-/-^* cells, but also dramatically increased (176-fold at 24 hpi) in *Casp1/11^-/-^* cells treated with α-IFNAR (**Fig. 4F**). We hypothesized that the protective effects of O-antigen against xenophagy may be revealed when blocking inflammasome and ISG activity, and therefore inhibited autophagy with the small molecule 3MA in these conditions; however, 3MA had only a minimal or no effect (**Supplemental Fig. 1B**), suggesting again that O-antigen did not play a major role in protecting *R. parkeri* from autophagy in macrophages, unlike OmpB. Together, these data suggest that O-antigen is critical for protecting *R. parkeri* from host inflammasome activation and IFN-β production.

We next asked if O-antigen protected against NO in the absence of inflammasome and interferon signaling. *Casp1/11^-/-^* BMDMs were treated with the α-IFNAR antibody as well as with the iNOS inhibitor L-NIL and infected with the O-antigen-deficient mutants. iNOS inhibition further increased bacterial burdens of *wecA* (37-fold at 72 hpi, **Fig. 4G**) and *rmlD* (15-fold at 72 hpi, **Fig. 4H**) mutants (**Fig. 4G**). iNOS inhibition did not significantly alter IFN-β or IL-1 β production during infection by O-antigen deficient *R. parkeri* in BMDMs (**Supplemental Fig. 3A-3D**). Thus, these studies revealed that the severe restriction of O-antigen-deficient mutants that we identified in the initial screen was largely due to killing by inflammasomes and ISGs, including GBPs and iNOS. This reveals a critical and multifaceted role for O-antigen in protecting *R. parkeri* from innate immunity in macrophages.

### *R. parkeri* O-antigen promotes cell-to-cell spread in epithelial cells independently of actin-based motility

We next examined whether *R. parkeri* O-antigen contributes to cell-to-cell spread, which is largely mediated by surface factors including RickA and Sca2 that promote actin-based motility. To determine if the O-antigen-deficient mutant bacteria had defects in spread, we measured the size of individual plaques formed upon infecting a confluent monolayer of Vero cells. This revealed that the *wecA* mutant had a plaque size of 47% relative to WT and the *rmlD* mutant had a plaque size of 67% relative to WT (**Supplemental Fig. 2A**). We hypothesized that O-antigen may affect actin-based motility and measured actin tails in WT and the O-antigen-deficient mutants. However, contrary to our hypothesis, actin tail formation was similar between WT bacteria and the mutants (**Supplemental Fig. 2B, C**). Together, these findings suggest that O-antigen promotes *R. parkeri* cell-to-cell spread in a mechanism independent of actin-based motility, although the explanations for this defect remain incompletely understood.

### O-antigen is essential for *R. parkeri* pathogenesis *in vivo*

We next sought to define the role for O-antigen in *R. parkeri* pathogenesis *in vivo.* Whereas WT C57bl/6 mice rapidly restrict *R. parkeri* and develop no or minor disease manifestations upon intradermal delivery, mice lacking the receptors for IFN-I (IFNAR) and IFN-γ (IFNGR) develop necrotic skin lesions (eschars) that are similar in gross pathology to those seen upon human *R. parkeri* infection^7^. These mice are a robust model system to study *R. parkeri* virulence genes, for example OmpB, PkmT1 and PkmT2 are required for *R. parkeri* to cause lethal disease in *Ifnar*^−/−^*Ifngr*^−/−^ mice^7,12^. However, the roles for WecA or RmlD in *R. parkeri* pathogenesis were unclear. We therefore intradermally infected *Ifnar^-/-^Ifngr^-/-^* mice with WT, *wecA,* and *rmlD* mutants and measured eschar formation and weight loss over time. Whereas WT *R. parkeri* caused severe eschars, the *rmlD* mutant caused an initial eschar that healed by day 20 (**Fig. 5A**). Most strikingly, the *wecA* mutant elicited no eschars and only mild redness throughout the time course (**Fig. 5A**). Whereas WT *R. parkeri* caused severe weight loss, suggesting systemic disease, the O-antigen-deficient mutants caused significantly less weight loss and remained above 95% of the original body weights (**Fig. 5B**). To mimic systemic disease, we next sought to determine if the O-antigen-deficient mutants caused weight loss or lethality upon intravenous delivery. Whereas the mice infected with WT *R. parkeri* succumbed, the mice infected with O-antigen-deficient mutants all survived (**Fig. 5C**) and had no significant weight loss (**Fig. 5D**). Together, these data indicate that O-antigen is a key determinant of *R. parkeri* pathogenesis *in vivo* that is required for causing eschars and disseminated disease.

**Figure 5:**
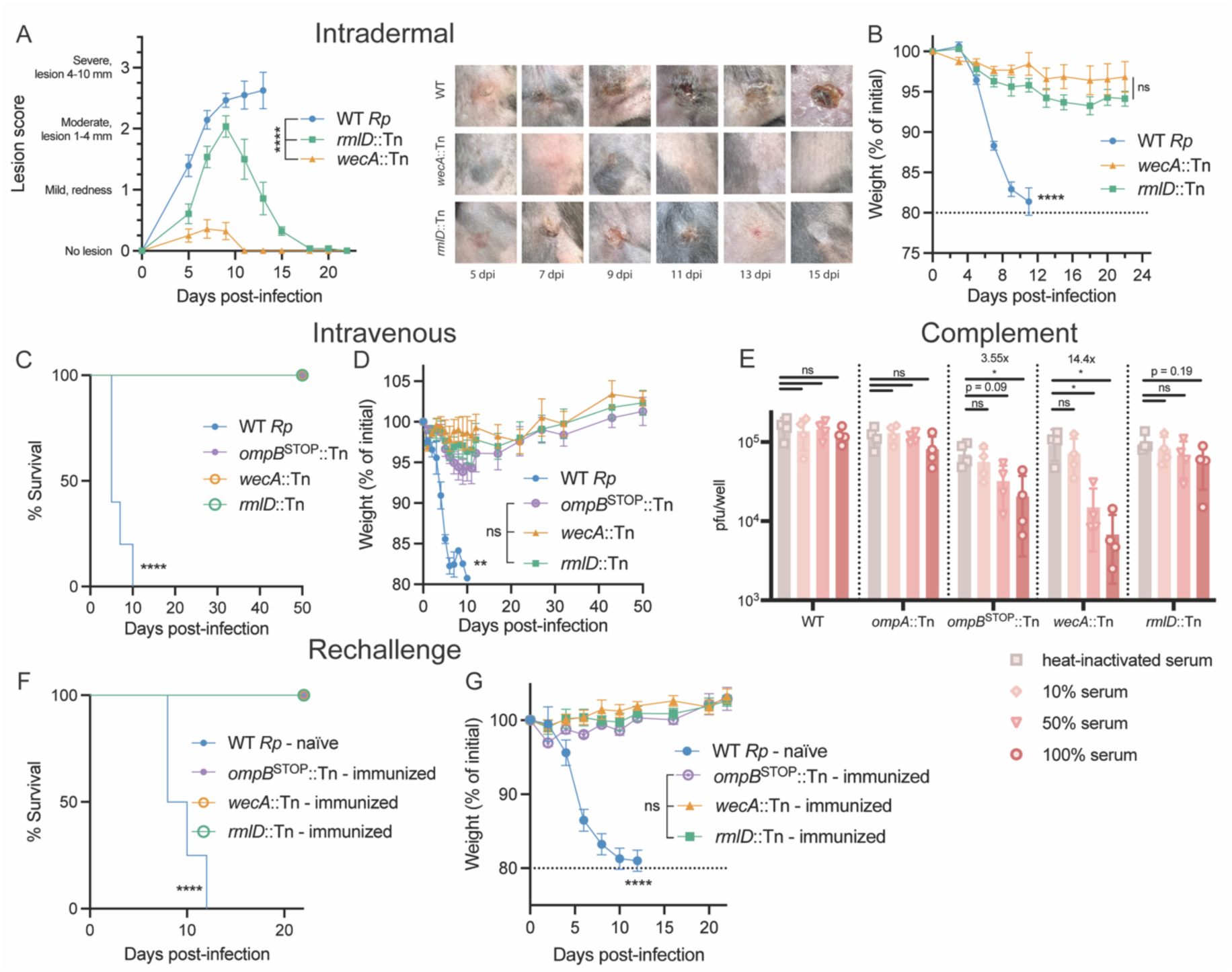
O-antigen is required for *R. parkeri* virulence *in vivo*. **A, B)** *Ifnar^-/-^Ifngr^-/-^* mice were intradermally infected with 10^5^ PFU of WT *R. parkeri* or indicated mutants and monitored over time for eschars and weight; WT n=7, *wecA*::Tn n=7, *rmlD*::Tn n=7. Statistical comparisons between WT and mutants were performed using a two-way ANOVA (left). Representative images of infected mice in (A) and (B) are shown for each bacterial strain. **B)** Weight of mice infected in (A). A mixed-effects model analysis from 0 to 25 days post-infection (dpi) was used to compare WT to mutants and a two-way ANOVA was used to compare between mutants from 0 to 25 dpi. **C, D)** *Ifnar^-/-^Ifngr^-/-^* mice were intravenously infected with 5 x 10^6^ PFU of the indicated *R. parkeri* strains; WT, n = 5; *wecA*::Tn, n = 6; *rmlD*::Tn, n = 6. Statistical comparison between WT and mutants was performed using a log-rank (Mantel-Cox) test. **D)** Weight changes of infected mice in (C). A mixed effects model analysis was used to compare WT and mutants from 0 to 50 days post-infection (dpi) and a two-way ANOVA from 0 to 50 dpi was used to compare mutants. **E)** 10^5^ PFU of the indicated *R. parkeri* mutants were incubated in concentrations of heat-inactivated and human serum (HS) from healthy blood donors for one hour at 33°C before being serially diluted onto confluent monolayers of Vero cells for plaque assay. Data are a combination of four independent experiments across four unique donors (n = 4). Statistics were calculated using a normal distribution one-way ANOVA and Brown-Forsyth and Welch ANOVA post-hoc comparisons. **F, G)** *Ifnar^-/-^Ifngr^-/-^*mice were intravenously infected with 5 x 10^6^ PFU of the indicated *R. parkeri* strains and then 40 dpi later rechallenged with 5 x 10^6^ PFU of WT *R. parkeri*. **F)** Survival of rechallenged mice infected with attenuated mutants (naïve - *Ifnar^-/-^Ifngr^-/-^*, n = 4; *wecA*::Tn – immunized, n = 6; *rmlD*::Tn – immunized, n = 6; *ompB^STOP^*::Tn – immunized, n = 5). **G)** Weight change of infected mice from (A). A mixed effects model analysis was used to compare WT and mutants from 0 to 50 days post-infection (dpi) and a two-way ANOVA from 0 to 50 dpi was used to compare mutants. All data are a combination of two independent experiments. All data are represented as mean ± SEM, * p < 0.05, **p<0.01, ****p < 0.0001, ns = not significant.

### O-antigen and OmpB promote *R. parkeri* resistance to complement

Complement proteins circulate in blood and can kill bacteria by assembling on their surface and disrupting membranes. Complement deficiency increases susceptibility to *R. australis*^32^ and *R. conorii* interacts with factor H and vitronectin to mediate complement resistance^33,34^. Antibodies to WT *R. conorii* promote killing by complement, while antibodies to O-antigen-deficient mutants do not^35^. Despite these advances, previous studies have not used mutant *Rickettsia* lacking O-antigen and OMPs to determine their roles in protection from complement. To investigate whether O-antigen and OMPs protect *R. parkeri* from complement, we incubated 10^5^ PFU of *wecA* and *rmlD* mutants in human sera, alongside WT bacteria as a control. We also included *R. parkeri* mutants lacking *ompA* and *ompB*, as these are also abundant surface factors which may mediate complement resistance^34^. Different concentrations of sera were heat-inactivated to ablate complement activity and PFU assays were performed to measure bacterial viability. These experiments revealed that *wecA* and *ompB* mutants were restricted 14-fold and 3-fold, respectively, in a manner that was dependent on the dose of viable sera, while the *rmlD* and *ompA* mutants were not (**Fig. 5E**). These data reveal that both O-antigen and OmpB play critical roles in mediating complement-mediated killing.

### Immunizing with O-antigen-deficient mutants protects against rechallenge *in vivo*

Understanding adaptive responses is critical for treating disease and vaccine design. Peptides from OMPs including OmpB can protect against infection via cell-mediated immunity^36–40^. In contrast, O-antigen is the target of antibodies^35^, which may not be as highly protective^38,41^. Previous studies have largely been performed without bacterial mutants lacking these factors, and we therefore sought to use the O-antigen-deficient mutants to elucidate the role for O-antigen in immunity. To this end, we immunized mice with *wecA* and *rmlD* mutant *R. parkeri* in *Ifnar^-/-^Ifngr^-/-^* mice and 40 days later rechallenged these mice with a lethal dose of WT bacteria. *ompB* mutants were used as a control, as we previously reported that they provided protection upon rechallenge^7^. All mice immunized with the O-antigen-deficient mutants survived, whereas all naïve mice succumbed to intravenous challenge by 12 d.p.i. (**Fig. 5F**). Upon intravenous rechallenge, mice immunized with *wecA*::Tn and *rmlD*::Tn also did not lose significant weight (**Fig. 5G**). As there are currently no available vaccines for rickettsiosis and considering that the *wecA* mutant elicited minimal disease upon intravenous or intradermal infection, this study reveals this strain as a potentially robust live attenuated vaccine platform.

### WecA and RmlD are required for the electron-lucent halo surrounding *R. parkeri*

We hypothesized that O-antigen served as a shield to protect the surface of *R. parkeri* from various innate immune assaults. *Rickettsia* species contain an electron-lucent halo surrounding the bacteria^42^, which is lost in *ompB* mutant *R. parkeri*^13^. We hypothesized that O-antigen contributed to halo formation and performed transmission electron microscopy (TEM) of WT, *wecA*::Tn, *rmlD*::Tn, and *ompB^stop^*::Tn bacteria. Similar to the *ompB* mutant bacteria, *wecA* and *rmlD* mutants also lost the halo (**Fig. 6A**). Together, these studies provide a model by which O-antigen protects against multiple innate immune defenses, including GBPs, nitric oxide, inflammasomes, complement, and E3 ligases (**Fig. 6B**).

**Figure 6:**
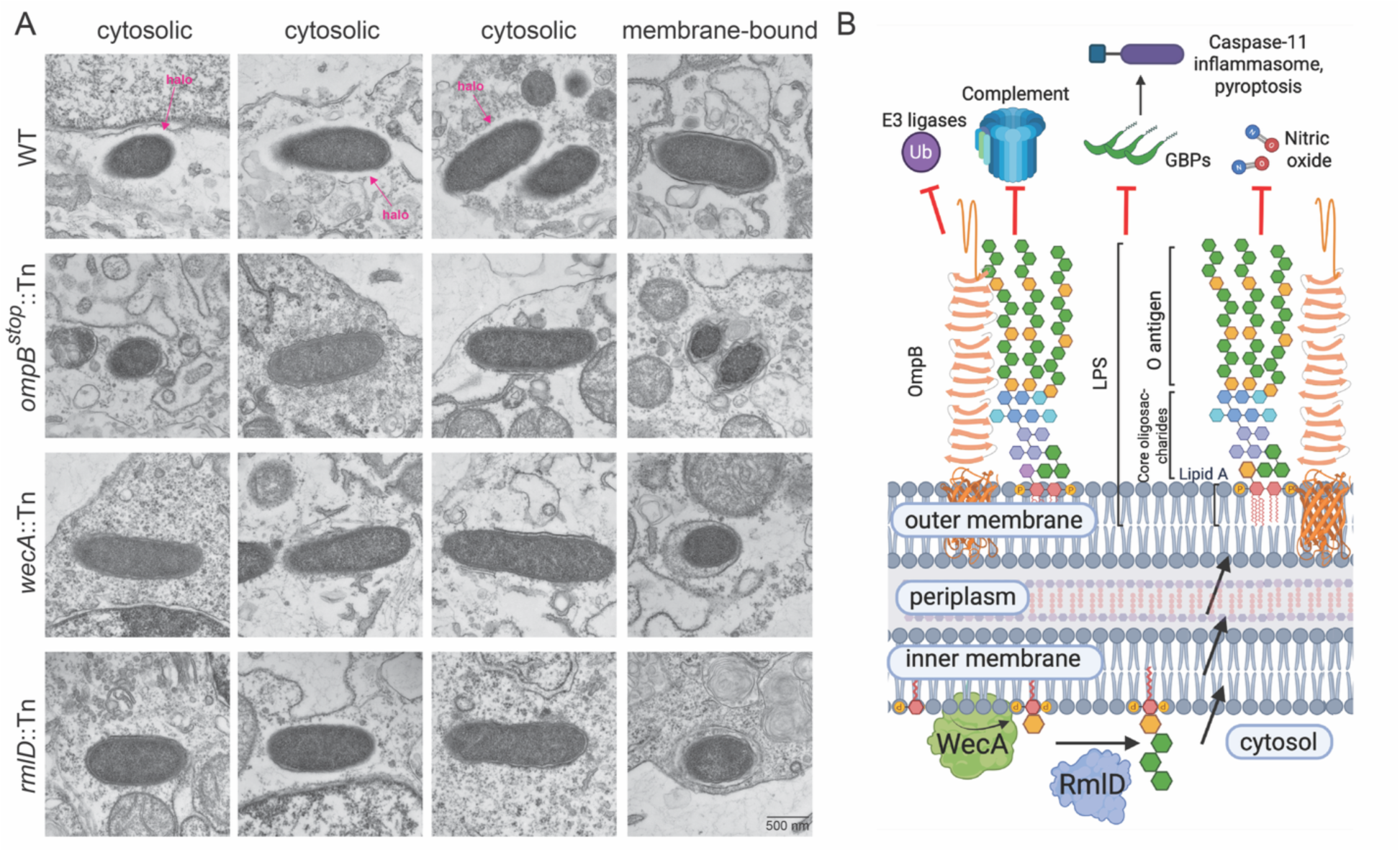
WecA and RmlD are required for the electron-lucent halo: **A)** TEM images of Vero cells infected with WT, *ompB^stop^*::Tn, *wecA*::Tn, and *rmlD*::Tn *Rp* in the indicated cellular locations at 1 hpi. Vero cells were infected at a MOI of 10. **B)** *R. parkeri* WecA and RmlD synthesize the first and second steps of O-antigen, which protects against GBP targeting, inflammasome activation, nitric oxide, complement and E3 ligases. Inflammasome activation was driven by both Caspase-1 and Caspase-11, although principally by Caspase-11. OmpB protects mainly from E3 ligases and xenophagy. The biophysical relationship between OmpB and O-antigen, for example which extends further from the bacterial surface and whether O-antigen penetrates the OmpB S-layer, remains unknown.

## Discussion

In this study we report the first forward genetic screen to identify factors required for *Rickettsia* species survival in macrophages, which are a primary cell type targeted *in vivo* by *Rickettsia* and many other intracellular bacterial pathogens^43,44^. This unbiased approach revealed WecA and RmlD, two factors critical for LPS O-antigen biosynthesis, as the most critical factors. LPS is highly immunostimulatory and it has remained unclear why it is conserved in obligate cytosolic pathogens. O-antigen was critical for protecting *R. parkeri* against inflammasomes, nitric oxide, and GBPs, altering the paradigm for how O-antigen protects cytosolic bacteria from GBPs. In mice, O-antigen was essential for virulence, including against complement, and these mutants elicited little disease while also eliciting long-lasting protection from rechallenge, thus providing promising live attenuated vaccine candidates. Through the lens of evolution, this study reveals why *Rickettsia,* as obligate intracellular pathogens, maintain immunostimulatory LPS, which is because it plays a multifaceted role in protecting from innate immunity.

As obligate intracellular microbes, evading innate immunity promotes *Rickettsia* longevity in host cells, where they survive for multiple days without causing host cell lysis. *R. parkeri* evades ubiquitin targeting and degradation by methylating the lysines on OMPs including OmpB with methyltransferases PkmT1 and PkmT2^12,13^. Thus, the major role for *R. parkeri* OmpB is to protect against xenophagy, whereas O-antigen is to protect against inflammasomes and ISGs. Although the *wecA* and *rmlD* mutants were ubiquitylated, albeit to a lesser extent than *ompB* mutants^12^, suggesting that the protective role against xenophagy is minor. We conclude that O-antigen and OmpB provide a complex and structured barrier against mammalian innate immunity (**Fig. 6B**). No secreted effectors have been reported in *Rickettsia* to directly modulate these eukaryotic innate immune factors, and thus *R. parkeri* appears to rely more on camouflaging its surface, contrasting other facultative intracellular pathogens like *Shigella flexneri,* which secretes at least nine effectors to modulate host cell death^45^. Future studies on the roles for *Rickettsia* surface proteins versus secreted factors in innate immune evasion will better shape our understanding of pathogenesis.

Both spotted fever group (SFG) and typhus group (TG) *Rickettsia* lipid A is hexa-acylated^46–48^ and immunostimulatory. For example, *Tlr4*^-/-^ mice are hypersusceptible to the SFG pathogen *Rickettsia conorii*^49^ and *Tlr4^-/-^*macrophages harbor increased burdens of *R. australis*^50^. Moreover, both SFG and TG *Rickettsia* species activate Caspase-11-dependent pyroptosis^19,20^. Since Caspase-11 detects the lipid A portion of LPS, similar to TLR4, this suggests that *Rickettsia* species do not modify their lipid A to evade these innate immune sensors. In contrast, *Francisella tularensis* has tetra-acylated lipid A that enables it to evade recognition by TLR4 and Caspase-11^21,51,52^. In another example, the obligate intracellular pathogen *Chlamydia trachomatis* has tetra-and penta-acylated lipid A, which only weakly activates TLR4^53^. Thus, it is an interesting contrast that certain pathogens like *F. tularensis* and *C. trachomatis* evolved non-canonical lipid A structures to avoid innate immunity, whereas *Rickettsia* species have not evolved such modifications. We speculate that perhaps there is a fitness cost associated with this LPS modification, and thus *Rickettsia* species instead protect against host innate immunity with O-antigen and OMPs. Our results here underscore the importance of *Rickettsia* O-antigen and not lipid A modifications in shielding against innate immunity.

This study reveals distinctions between *R. parkeri* and other intracellular pathogens in their interactions with innate immunity. In contrast to our findings with *R. parkeri*, *F. tularensis* O-antigen enables autophagic evasion, whereby O-antigen-deficient mutants are ubiquitylated and restricted by xenophagy^54^. *F. tularensis* O-antigen also promotes survival in macrophages by protecting against AIM2 inflammasome-mediated cell death^31,55^, whereas we found no role for AIM2. Although GBPs can protect against *F. novicida*^56^, the role for *Francisella* O-antigen in protecting against GBPs is unknown. Another contrasting finding to our studies on *R. parkeri* is that *Shigella flexneri* O-antigen-deficient mutants have decreased targeting by GBPs^28^, suggesting that *S. flexneri* O-antigen actually recruits GBPs. This established a model that GBPs may target O-antigen^27^, and thus our findings with *R. parkeri* represent a novel paradigm for bacterial defense against GBPs, whereby O-antigen shields against GBP-mediated recognition and subsequent inflammasome activation.

The importance of *R. parkeri* O-antigen for intracellular survival in macrophages largely aligns with findings on O-antigen in the SFG pathogen *R. conorii,* where O-antigen contributes to plaque formation and invasion *in vitro*^35^. Notably, among the two O-antigen-deficient mutants identified in *R. conorii,* both grew in BMDMs^57^, whereas we found that the *wecA* and *rmlD* mutants in *R. parkeri* were severely attenuated. One of the *R. conorii* O-antigen-deficient mutants HK2, is attenuated for virulence *in vivo*, while the other still caused lethal disease^35^. HK2 protected from rechallenge^57^, similar to our findings with *wecA* and *rmlD*. The *R. conorii* O-antigen-deficient mutants may retain the electron-lucent halo, which we speculate is because these mutants still retain some polysaccharide synthesis, whereas WecA and RmlD likely do not, as they are required for the first and second steps of O-antigen. Our study builds on the previously reported findings by revealing the mechanisms of restriction in macrophages through inflammasomes, GBPs, and nitric oxide and by determining an additional mechanism of restriction *in vivo* by revealing that the mutants are more susceptible to complement. As our initial screen was an unbiased comparison to >350 other *R. parkeri* mutants, our study also reveals O-antigen as among the most critical virulence factors to date, along with OmpB, for *R. parkeri* virulence. As our screen was not saturating, additional screening may identify other critical virulence factors that promote survival in macrophages.

Understanding adaptive responses is critical for treating disease and vaccine design. Peptide vaccines from OMPs including OmpB can protect against infection via cell-mediated immunity^36–40^. In contrast, rickettsial O-antigen is the target of antibodies^35^, which may not be as highly protective^38,41^. Due to a historical lack of genetic tools and mutants, many previous studies on adaptive responses to *Rickettsia* did not use mutant bacteria. Our study demonstrates that robust protective responses to WT can be generated by O-antigen-deficient bacteria, suggesting that other surface antigens besides O-antigen can be strongly protective.

In summary, we report that rickettsial O-antigen is critical for protecting against innate immunity by mechanisms that are distinct from other intracellular Gram-negative bacteria, and this knowledge has translational importance for vaccine design. Lingering questions in the field are why OmpB protects against xenophagy while O-antigen protects against inflammasomes and ISGs, and how these entities interact with one another physically, for example whether O-antigen penetrates the OmpB S-layer. A second unanswered question is why these mutants have small plaques in Vero cells despite growing at similar rates. A third unanswered question is why *wecA* and *rmlD* mutant growth is not completely restored to WT levels in BMDMs lacking inflammatory caspases, interferon signaling, and nitric oxide. Answering this question may reveal additional layers of innate immunity that protect macrophages but not epithelial cells from intracellular invaders. Lastly, regarding immunity, it is unclear whether the *R. parkeri* or *R. conorii* O-antigen-deficient mutants broadly protect against different SFG *Rickettsia* species, for example *R. rickettsii,* which would provide valuable insight into their potential as vaccine candidates. Answering these questions will enhance our understanding of how O-antigen plays such a crucial role in *Rickettsia* species intracellular survival in immune cells.

## Methods

### Preparing R. parkeri

*R. parkeri* strain Portsmouth was originally obtained from Christopher Paddock (Centers for Disease Control and Prevention) who provided it to Dr. Matt Welch (UC Berkeley). Bacteria were amplified in confluent T175 flasks of Vero cells (female African green monkey kidney epithelial cells), which were obtained from the UC Berkeley Cell Culture Facility, where they were tested for mycoplasma contamination, and were authenticated by mass spectrometry experiments. Vero cells were grown in DMEM (Gibco 11965-092) with glucose (4.5 g/L) supplemented with 2% fetal bovine serum (FBS, Corning 35-010-CV). To prepare bacteria for infection, T175 flasks of confluent Vero cells were infected with 5 x 10^6^ *R. parkeri* per flask and at 5 dpi when cells were heavily infected, infected cells were scraped, collected, and centrifuged at 12,000 x G for 20 min at 4°C. Pelleted cells were then resuspended in K-36 buffer (0.05 M KH_2_PO_4_, 0.05 M K_2_HPO_4_, 100 mM KCl, 15 mM NaCl, pH 7) and dounced (60 strokes) on ice. The solution was then centrifuged at 200 x G for 5 min at 4°C to pellet host cell debris. Supernatant containing *R. parkeri* was overlaid on a 30% Ficoll (Cytiva, 17144002) solution in K-36 buffer. Gradients were centrifuged at 18,000 rpm in an SW-28 ultracentrifuge swinging bucket rotor (Beckman/Coulter) for 20 min at 4°C to separate host cell debris. Bacterial pellets were resuspended in brain heart infusion (BHI) media (BD, 237500) and stored at -80°C. Bacterial titers were determined via plaque assays by serially diluting the bacteria in 12-well plates containing confluent Vero cells. Plates were then spun for 5 min at 300 x G in an Eppendorf 5810R centrifuge. Titers were determined using the plaque assay described below.

### Genetic screen in BMDMs

Mutants that were screened for survival in BMDMs were previously generated by electroporation with the Himar1 transposon pMW1650^12,22^. 350 total mutants were screened, whereby each mutant was thawed and diluted into two wells of a 24 well of BMDMs containing 10^5^ BMDMs, plated the day before. At 2 and 72 hpi, one well for each mutant were lysed with water and plated onto confluent Vero cells in 12 well plates. If any mutant strain had greater than 10^4^ PFU at 2 hpi, that strain was repeated with a lower dilution. If any mutant gave zero plaques at 2 hpi, that strain was repeated with an increased dilution from the stock. WT and *ompB^stop^*::Tn bacteria (this mutant is described Engström et al., 2019) were used in each experiment as controls. The identity of all other samples was blinded. Hits for further analysis were those with a growth rate below 1 and were retested using more timepoints from 24-72 hpi. The genomic insertion sites of the genes of interest were: *wecA*::Tn (1223170) and *rmlD* (455753).

### Deriving bone marrow macrophages

Animal research using mice was conducted under a protocol approved by the UC Irvine Institutional Animal Care and Use Committee (IACUC) in compliance with the Animal Welfare Act and other federal statutes relating to animals and experiments using animals (Burke lab animal use protocol AUP-25-004). The UC Irvine IACUC is fully accredited by the Association for the Assessment and Accreditation of Laboratory Animal Care International and adheres to the principles of the Guide for the Care and Use of Laboratory Animals. To obtain bone marrow, male or female mice were euthanized, and femurs, tibias, and fibulas were excised. Connective tissue was removed, and the bones were sterilized with 70% ethanol. Bones were washed with BMDM media (20% Corning FBS, 1% sodium pyruvate, 0.1% β-mercaptoethanol, 10% conditioned supernatant from 3T3 fibroblasts, in Gibco DMEM containing glucose and 100 U/ml penicillin and 100 ug/ml streptomycin) and ground using a mortar and pestle. Bone homogenate was passed through a 70 μm nylon Corning Falcon cell strainer (Falcon 352350) to remove particulates. Filtrates were centrifuged in an Eppendorf 5810R at 1,200 RPM (290 x G) for 8 min, supernatant was aspirated, and the remaining pellet was resuspended in BMDM media. Cells were then plated in non-TC-treated 15 cm petri dishes (at a ratio of 10 dishes per 2 femurs/tibias) in 30 ml BMDM media and incubated at 37° C. An additional 30 ml media without antibiotics was added 3 d later. At 7 d the media was aspirated, and cells were incubated at 4°C with 15 ml cold PBS (Gibco, 10010-023) for 10 min. BMDMs were then scraped from the plate, collected in a 50 ml conical tube, and centrifuged at 1,200 RPM (290 x G) for 5 min. The PBS was then aspirated, and cells were resuspended in BMDM media with 30% FBS and 10% DMSO at 1.2×10^7^ cells/ml. 1 ml aliquots were stored in liquid nitrogen.

### Serum collection

Healthy blood donors were collected at the UC Irvine Institute for Clinical and Translational Science (ICTS) and allowed to clot for 20 minutes at room temperature before centrifugation at 2,000 RCF at 4°C for 10 minutes. Donor sera were individually aliquoted for storage at -80 °C and used within 6 months of collection.

### *In vitro* experiments

To plate cells for infection, aliquots of BMDMs were thawed on ice, diluted into 9 ml of DMEM, centrifuged in an Eppendorf 5810R at 1,200 RPM (290 x G) for 5 minutes, and the pellet was resuspended in 10 ml BMDM media without antibiotics. The number of cells was counted using Trypan blue (Sigma, T8154) and a hemocytometer (Bright-Line), and 5 x 10^5^ cells were plated into 24-well plates. Approximately 16 h later, 30% prep *R. parkeri* were thawed on ice and diluted into fresh BMDM media to the desired concentration. Media was then aspirated from the BMDMs, replaced with 500 µl media containing *R. parkeri,* and plates were spun at 300 G for 5 min in an Eppendorf 5810R. Infected cells were then incubated in a humidified CEDCO 1600 incubator set to 33°C and 5% CO_2_ for the duration of the experiment.

Titers were determined via plaque assays by serially diluting the bacteria in 12-well plates containing confluent Vero cells. Plates were then spun for 5 min at 300 x G in an Eppendorf 5810R centrifuge. At 24 hpi, the media from each well was aspirated and the wells were overlaid with 2 mL/well DMEM with 5% FBS and 1.2% Avicel^®^ PH-101 (Sigma,11363). At 7 dpi, the wells were fixed with 2 mL/well of 7% Formaldehyde (VWR,10790-708) for 30 minutes and with 0.5 mL/well of 1x crystal violet (VWR, 470337-534) for 15 minutes. Crystal violet was then washed off with tap water and plaques were counted.

For experiments involving α-IFNAR antibody, 5 x 10^5^ of respective BMDMs were plated per well in 24 well plates. The following day, media was aspirated, cells were infected with *R. parkeri* at a MOI of 1. 1 mM α-IFNAR-1 (mouse, Leinco Technologies, I-401) was added to each well 1 hour postinfection. BMDMs were then lysed at different timepoints and plated on Vero cells for quantification via PFU assay.

For experiments involving L-NIL, 5 x 10^5^ of *Casp 1/11^-/-^* BMDMs were plated per well in 24 well plates. The following day, media was aspirated, cells were infected with *R. parkeri* at a MOI of 1. 1 mM L-NIL (MedChemExpress, HY-12116) was added to each well 1 hour postinfection. BMDMs were then lysed at different timepoints and plated on Vero cells for quantification via PFU assay.

For experiments with NONOate, the compounds were resuspended to 150 mM in MilliQ water, stored at −80°C in single-use aliquots, and directly added to *R. parkeri* containing cultures. For direct incubations, *R. parkeri* was incubated in brain heart infusion serum (BHI, BD Difco™, DF0037178) for 30 minutes with indicated amounts of DEA-NONOate (Cayman Chemicals, 82100). For experiments involving ATP, *R. parkeri* was incubated in BHI for 2 hours with indicated amounts of Spermine-NONOate (Cayman Chemicals, 82150) along with the addition of 1 mM ATP. Incubations occur in non-TC treated 96 well plates at 33°C and 5% CO_2_. At indicated timepoints, *R. parkeri* was then plated on Vero cells for quantification via PFU assay.

For experiments with autophagy inhibitors, 5 x 10^5^ of WT and *Casp 1/11^-/-^* BMDMs were plated per well in 24 well plates. The following day, media was aspirated, cells were infected with *R. parkeri* at a MOI of 1. 5 mM 3-methyladenine (3MA, Selleck, S2767) was added to each well one hour postinfection. BMDMs were then lysed at different timepoints and plated on Vero cells for quantification via PFU assay.

For LDH assays, 50 µl of supernatant from wells containing BMDMs was collected into 96-well plates and LDH assay buffer was added according to manufacturer’s instructions (Promega CytoTox 96 Non-Radioactive Cytotoxicity Assay, G1780). Supernatant from uninfected cells was used as a negative control and from cells lysed with 1% triton X-100 (final concentration) was a positive control. Reactions were incubated at room temperature for 30 min prior to reading at 490 nm using a ClarioStar-Plus plate reader. Values for uninfected cells were subtracted from the experimental values, divided by the difference of triton-lysed and uninfected cells, and multiplied by 100 to obtain percent lysis. Each experiment was performed and averaged between technical duplicates and biological triplicates.

For the IL-1β (R&D Systems, DY401) and IFN-b (R&D Systems, DY823405) ELISA, supernatants were collected at 24 hours postinfection from WT, *Gbp^chr3-/-^*, *Casp 1/11^-/-^, Casp 1^-/-^, Casp 11^-/-^, AIM2^-/-^,* and *NLRP3^-/-^* BMDMs infected at a MOI of 1 of *R. parkeri.* The ELISA was done according to manufacturer’s instructions and read using a ClarioStar-Plus plate reader at 450 nm.

For assessing bacterial survival against complement, serum designated for heat-inactivation was thawed at room-temperature before being submerged in a water bath maintained at ∼56°C for 30 minutes and swirled every 10 minutes. Normal human serum was thawed at 4°C before being maintained on ice. *R. parkeri* aliquots between 2-6 mL of BHI were incubated in 200 mL of human serum in non-TC treated 96-well plates for 1 hour at 33°C before enumerating viable bacteria through plaque assay.

### Immunofluorescence microscopy

For immunofluorescence microscopy, 5 x 10^5^ HMEC-1 (originally obtained from Dr. Matt Welch, UC Berkeley), or BMDMs were plated overnight in 24-well plates on sterile 12 mm coverslips (Thermo Fisher Scientific, 12-545-80). Infections were performed as described above. At the indicated times post-infection, coverslips were washed once with PBS and fixed in 4% paraformaldehyde (Ted Pella Inc., 18505, diluted in 1 x PBS) for 10 min at room temperature. Coverslips were then washed 3 times in PBS. Coverslips were washed once in blocking buffer (1 x PBS with 2% BSA) and permeabilized with 0.5% triton X-100 for 10 min. Coverslips were incubated with primary or secondary antibodies diluted in 2% BSA in PBS for 30 min at room temperature. *R. parkeri* was detected using anti-*Rickettsia* 14-13 (mouse) diluted 1:250. Nuclei were stained with DAPI (BD Biosciences 564907) diluted 1:2,000, GBP2 was stained with a GBP2 rabbit antibody (Proteintech, 118541AP150UL) diluted 1:50, and actin was stained with Alexa-568 phalloidin (Life Technologies, A12380) diluted 1:500. Secondary antibodies were AlexaFluor-488 goat anti-mouse (Invitrogen A11001, lot 2513496), AlexaFluor-647 goat anti-rabbit (Invitrogen A32733, lot XG349344), all diluted 1:500. Coverslips were mounted in ProLong™ Gold Antifade Mountant (Invitrogen, P36934). Samples were imaged with either the Keyence BZ-X810 Inverted Microscope with a 60x oil objective or the Olympus FV3000 Laser-Scanning Confocal Spectral Inverted Microscope with Plan-Apochromat 60x/1.42 Oil (FWD=0.15mm) objective at the UC Irvine Microscope Imaging Core. Images were processed using (ImageJ) FIJI^49^ version 2.16.0/1.54p and brightness and contrast adjustments were applied to entire images. Images were assembled using Adobe Illustrator. Representative images are a single optical section. For quantification and cell length measurements, images were blinded. Images were analyzed using FIJI, descriptions of the number of bacteria counted are described in figure legends. Generally, at least 300 bacteria, consisting of combining at least three separate images per sample per time point per experiment. Bacteria were counted if they were in the plane of view and no bacteria in each plane were excluded.

### Transmission Electron Microscopy (TEM)

For the TEM experiments, Vero cells were infected at a MOI of 10. For ultrastructural analysis of cytoplasmic bacteria, infected cells were fixed in 2% paraformaldehyde/2.5% glutaraldehyde (Ted Pella Inc., Redding, CA) in 100 mM cacodylate buffer, pH 7.2 for 2 hr at room temperature. Samples were washed in cacodylate buffer and postfixed in 1% osmium tetroxide (Ted Pella Inc.) 1.5% potassium ferricyanide (Sigma, St. Louis, MO) for 1 hr. Samples were then rinsed extensively in dH20 prior to en bloc staining with 1% aqueous uranyl acetate (Ted Pella Inc.) for 1 hr. Following several rinses in dH20, samples were dehydrated in a graded series of ethanol and embedded in Eponate 12 resin (Ted Pella Inc.). Ultrathin sections of 95 nm were cut with a Leica Ultracut UCT ultramicrotome (Leica Microsystems Inc., Bannockburn, IL), stained with uranyl acetate and lead citrate, and viewed on a JEOL 1200 EX transmission electron microscope (JEOL USA Inc., Peabody, MA) equipped with an AMT 8 megapixel digital camera and AMT Image Capture Engine V602 software (Advanced Microscopy Techniques, Woburn, MA).

### In Vivo Experiments

*Ifnar^-/-^Ifngr^-/-^* mice were purchased from Jackson Labs (*Ifnar1*^-/-^*Ifngr1*^-/-^), strain #029098. All mice were healthy at the time of infection and were housed in microisolator cages and provided chow, water, and bedding. No mice were administered antibiotics or maintained on water with antibiotics. Experimental groups were littermates of the same sex that were randomly assigned to experimental groups, and each group for every experiment contained similar numbers of both male and female mice. Mice selected for experiments were age matched between groups and were between 8 and 35 weeks old at the time of initial infection. After the first experiment, a Power Analysis was conducted to determine subsequent group sizes. The sex of mice used for survival after infection and raw data for mouse experiments are provided in the Source Data for each figure. All mice were of the C57BL/6J background. For mouse infections, *R. parkeri* was prepared by diluting 30%-prep bacteria into cold sterile PBS on ice, centrifuged to pellet the bacteria, and resuspended in PBS. Bacterial suspensions were kept on ice during injections. For i.d. infections, mice were anaesthetized with 2.5% isoflurane via inhalation. The right flank of each mouse was shaved with a hair trimmer (Braintree CLP-41590), wiped with 70% ethanol, and 50 µl of bacterial suspension in PBS was injected intradermally using a 29-gauge needle. Mice were monitored for ∼3 min until they were fully awake. No adverse effects were recorded from anesthesia. For i.v. infections, mice were exposed to a heat lamp while in their cages for approximately 5 min and then each mouse was moved to a mouse restrainer (Braintree, TB-150 STD). The tail was sterilized with 70% ethanol, and 200 µl of bacterial suspension in sterile PBS was injected using 29-gauge needles into a lateral tail vein. All mice in this study were monitored daily for clinical signs of disease throughout infection, such as hunched posture, lethargy, scuffed fur, paralysis, facial edema, and lesions on the skin of the flank and tail. Mice were also monitored daily for changes in body weight. If a mouse displayed severe signs of infection, as defined by lethargy that prevented normal movement, body temperature below 90 °F, or a weight loss of ≤20% their starting weight, the animal was immediately and humanely euthanized using CO_2_ followed by cervical dislocation, according to IACUC-approved procedures. Pictures of skin lesions were obtained with permission from the Animal Care and Use Committee Chair and the Office of Laboratory and Animal Care. Pictures were captured with an Apple iPhone 11 Pro Max and images were evenly exposed in Apple Preview. All data are combined from at least two independent experiments.

### Structural predictions

Alphafold 3 was used for structural predictions of *R. parkeri* protein WecA (MC1_07070) and RmlD (MC1_02580) which provided predicted template modeling (pTM) scores of 0.95 and 0.96, respectively. The structural predictions were searched in FoldSeek 10 and the highest similarity to a homolog outside of the *Rickettsia* species for WecA was from the AFDB-PROTEOME database (glycosyltransferase WbpL from Pseudomonas aeruginosa PAO1, TM-score 0.847, RMSD: 3.56, E-Value 1.63e-16). For RmlD the homolog with the highest similarity was from the AFDB50 database (dTDP-4-dehydrorhamnose reductase from *Arcobacter* Sp. CECT 8986, TM-score: 0.976, RMSD: 0.96, E-value 2.54e-43).

### Statistical analysis

Statistical parameters and significance are reported in the figure legends. For comparing two sets of data, a two-tailed Student’s T test was performed. For comparing multiple data sets, a one-way ANOVA with multiple comparisons with Tukey post-hoc test was used for normal distributions, and a Mann-Whitney *U* test was used for non-normal distributions. A two-way ANOVA was used for data with a second variable. Lognormal tests were used when the logarithmic data was normally distributed. For *in vivo* experiments, sample sizes were based on previous studies of *R. parkeri* infection and were sufficient to detect biologically meaningful differences. Data were determined to be statistically significant when p<0.05. Asterisks denote statistical significance as: *, p<0.05; **, p<0.01; ***, p<0.001; ****, p<0.0001, compared to indicated controls. Error bars indicate standard deviation (SD). All other graphical representations are described in the Figure legends. Statistical analyses were performed using GraphPad PRISM.

## Data availability

*R. parkeri* strains were authenticated by whole genome sequencing and are available in the NCBI Trace and Short-Read Archive; Sequence Read Archive (SRA), accession number SRX4401164. Source data are provided in the Dryad repository. No original code was developed in this work.

## Acknowledgements

T.P.B. was supported by the National Institute of Allergy and Infectious Diseases (NIAID) of the National Institutes of Health (NIH) under Award Number R01AI185119. A.A.G. was supported by NIH-MARC U-STAR training grant T34GM136498. The content is solely the responsibility of the authors and does not necessarily represent the official views of the NIH. Models were made using BioRender. WT *R. parkeri* and all transposon mutants were a generous gift from Dr. Matthew Welch (University of California, Berkeley). *Atg5*^-/-^ and *Atg5^fl^*^/fl^ cells were a generous gift from Dr. Christina Stallings (Washington University). *Gbp^chr^*^3-/-^ cells were a generous gift from Dr. Jörn Coers (Duke University). Immortalized BMDMs (L3OG) were a generous gift from Dr. Andrew Olive (Michigan State University).

## Additional Information

Correspondence and requests for materials should be addressed to T.P.B.

## Competing interests

The authors declare no competing interests.

## Notes

### Competing Interest Statement

The authors have declared no competing interest.

