## Supplemental Figures for "A genetic screen identifies O-antigen as essential for *Rickettsia parkeri* survival in macrophages by protecting from inflammasomes and interferon-stimulated genes"

**Sun et al., 2026**

### Supplementary Figures:

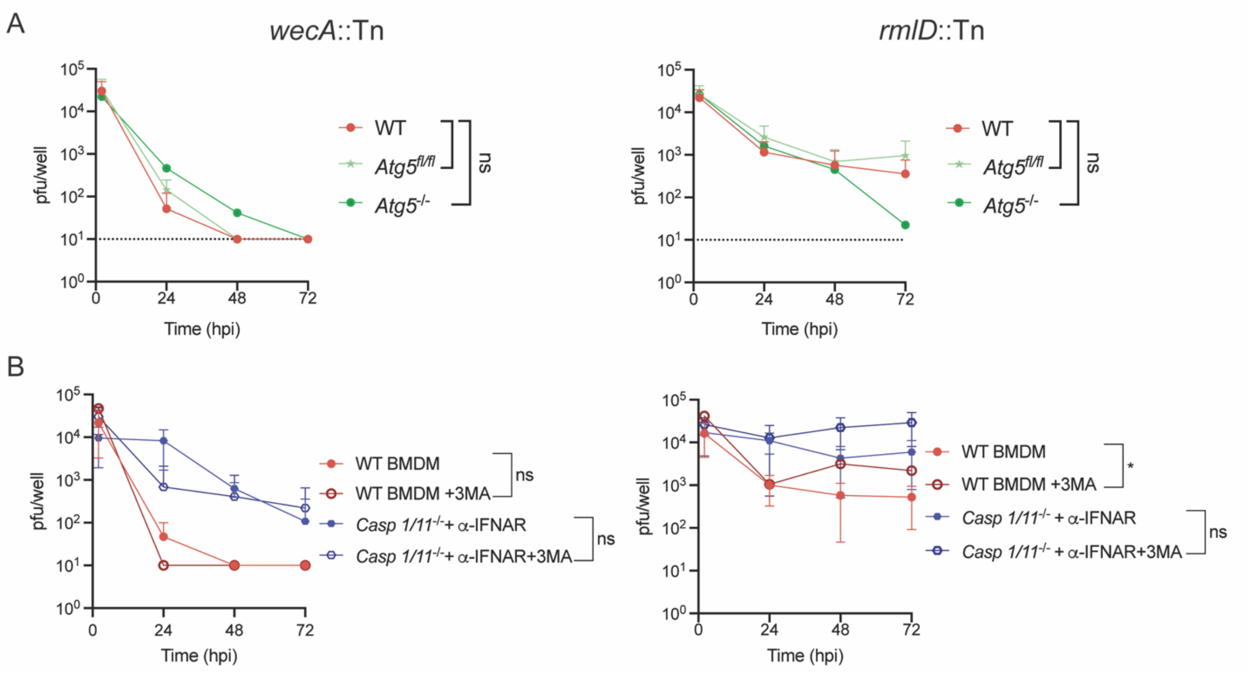


**Figure S1: Autophagy is not a major host defense restricting O-antigen mutants in macrophages. A)** Abundance of *wecA*::Tn and *rmlD*::Tn (n = 8) in WT, *Atg5$^{fl/fl}$, and Atg5$^{-/-}$* BMDMs infected at an MOI of 1. **B)** Abundance of *wecA*::Tn and *rmlD*::Tn (n = 8) in WT and *Casp 1/11$^{-/-}$* BMDMs with the addition of anti-IFNAR antibody and 3MA at indicated conditions. BMDMs were infected at an MOI of 1.1mM Anti-IFNAR antibody and 1mM 3MA were added one-hour postinfection. Infected cells were lysed at indicated timepoints for plaque assay. Dotted line indicates the limit of detection. Data are expressed as means ± SD. Statistics were performed with a two-way ANOVA, *$p<0.05$, **$p<0.01$, ns = not significant.

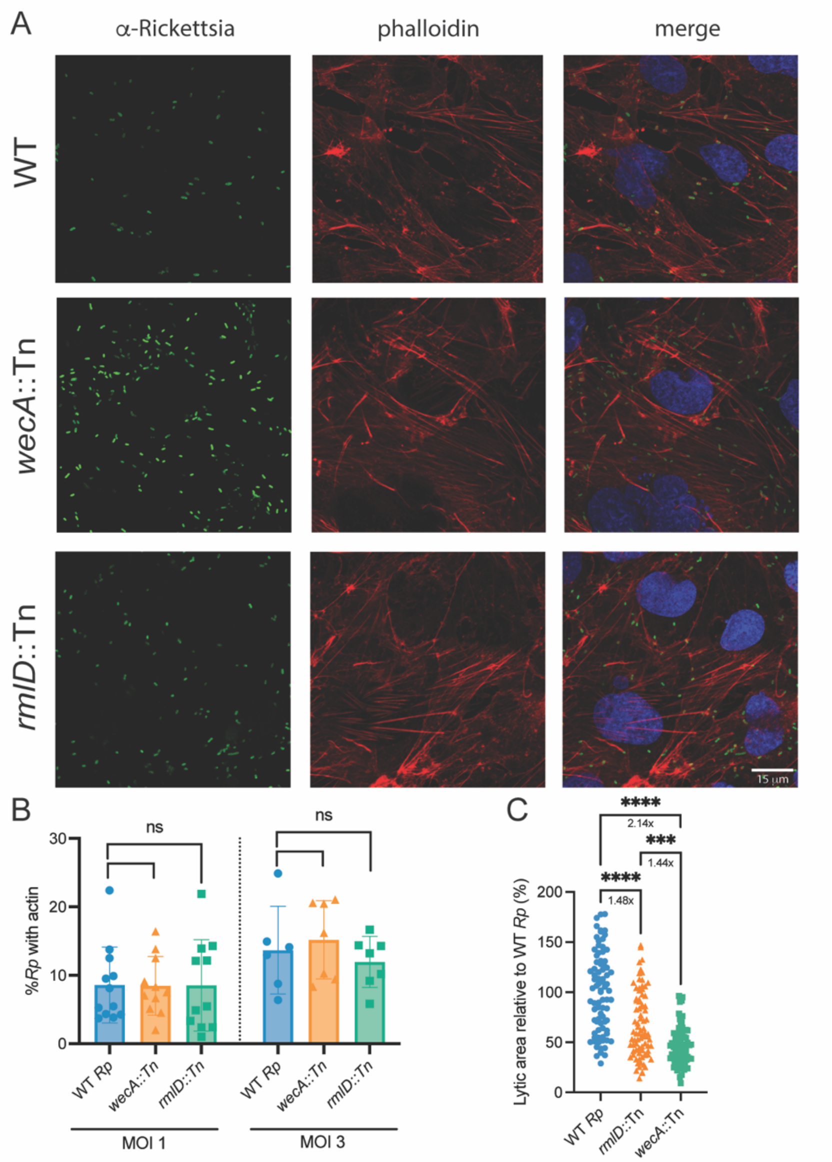


**Figure S2: *R. parkeri* O-antigen promotes cell-to-cell spread in epithelial cells but does not affect actin-based motility.**

**A)** Representative images of HMECs infected with WT, *wecA*::Tn and *rmlD*::Tn *R. parkeri* at an MOI of 1 and imaged at 48 hpi. Coverslips were stained with a phalloidin antibody to visualize actin (red) and an anti-Rickettsia antibody (green). Scale bar = 15 µm. **B)** Quantification of actin colocalization during WT, *wecA*::Tn and *rmlD*::Tn *R. parkeri* infection, from images in **(A)**. Data was a compilation of 3 separate experiments and a total of >300 bacteria were quantified per condition. Statistics were performed with a One-Way ANOVA. **C)** Quantification of plaque area of *wecA*::Tn and *rmlD*::Tn relative to WT *R. parkeri* (n = 83). Data are expressed as means ± SD and were compared with a One-way ANOVA, ***p<0.001, ****p<0.0001, ns = not significant.

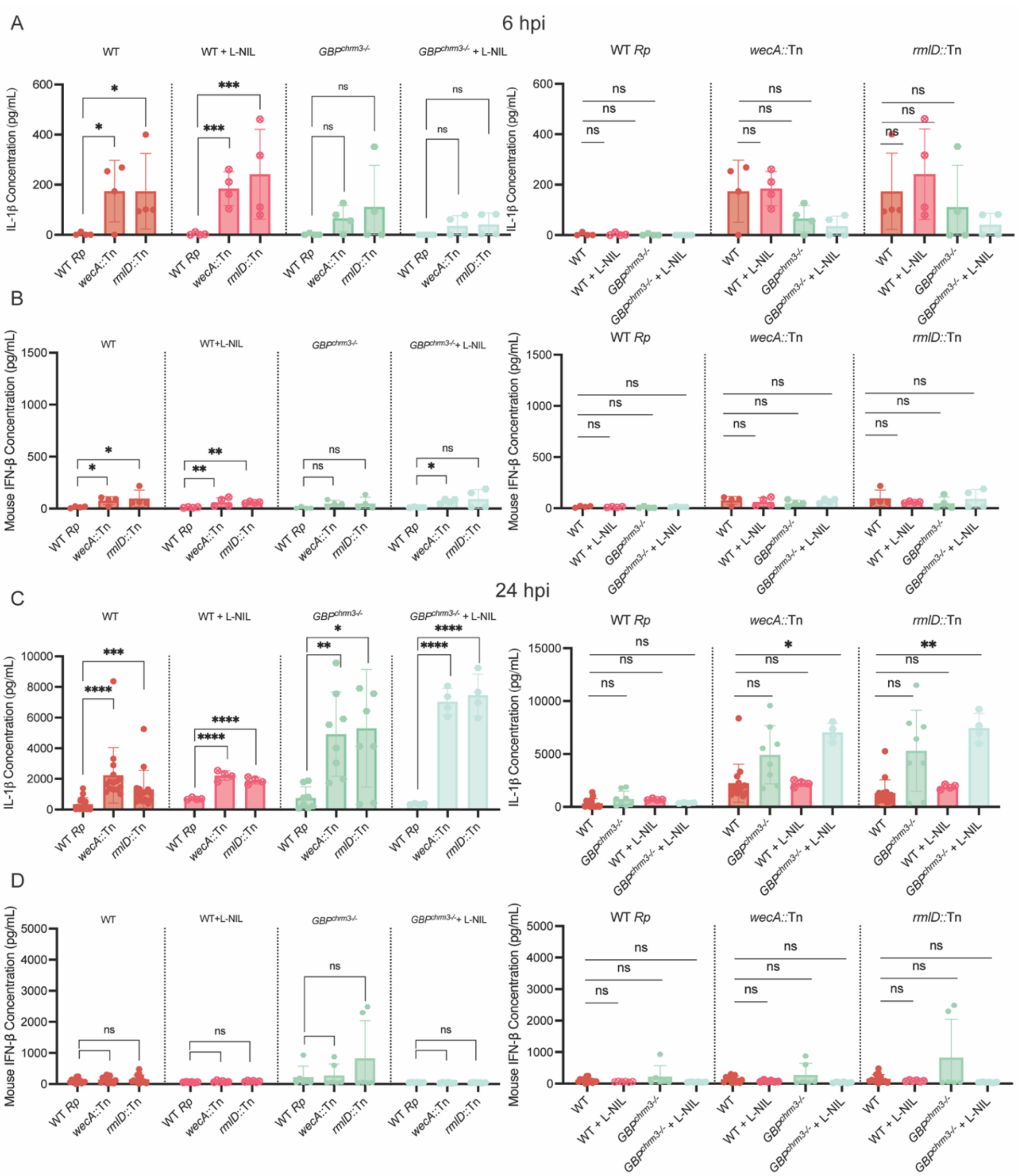


**Figure S3: iNOS inhibition does not significantly alter IFN-β or IL-1β production during infection by O-antigen deficient *R. parkeri*.**

**A)** Quantification of IL-1β concentration during WT, *wecA*::Tn and *rmlD*::Tn *R. parkeri* infection of WT and *GBP*$^{chrm3-/-}$ BMDMs with or without the addition of L-NIL (n=4). IL-1β concentration was measured at 6 hpi after *R. parkeri* infection at an MOI of 1. **B)** Quantification of IFN- β concentration during WT, *wecA*::Tn and *rmlD*::Tn *R. parkeri* infection of WT and *GBP*$^{chrm3-/-}$ BMDMs with or without the addition of L-NIL (n=4). IFN-β concentration was measured at 6hpi after *R. parkeri* infection at an MOI of 1. **C)** Quantification of IL-1β concentration during WT, *wecA*::Tn and *rmlD*::Tn *R. parkeri* infection of WT BMDMs (n=15), *GBP*$^{chrm3-/-}$ BMDMs (n=8), WT BMDMs + L-NIL (n=4) and *GBP*$^{chrm3-/-}$ BMDMs + L-NIL (n=4) at 24 hours postinfection. **D)** Quantification of IFN-β concentration during WT, *wecA*::Tn and *rmlD*::Tn *R. parkeri* infection of WT BMDMs (n=13), *GBP*$^{chrm3-/-}$ BMDMs (n=6), WT BMDMs + L-NIL (n=6) and *GBP*$^{chrm3-/-}$ BMDMs + L-NIL (n=6) at 24 hours postinfection. 1mM L-NIL was added at indicated conditions one-hour postinfection. Data are expressed as means ± SD. Statistics were performed with a Lognormal One-Way ANOVA, $*p<0.05$, $**p<0.01$, $***p<0.001$, $****p<0.0001$, ns = not significant.
